# Dynamics and interactions of ADP/ATP transporter AAC3 in DPC detergent are not functionally relevant

**DOI:** 10.1101/317669

**Authors:** Vilius Kurauskas, Audrey Hessel, François Dehez, Chris Chipot, Beate Bersch, Paul Schanda

## Abstract

A recent study^1^ used solution-state NMR spectroscopy to examine the interactions and dynamics of the yeast mitochondrial inner-membrane ADP/ATP carrier, yAAC3. Crystal structures of different AACs, including yAAC3, in a conformation locked with a strong inhibitor (CATR) had been determined before. This putative “c-state”^2,3^ is believed to represent one extreme conformation of an alternating access mechanism, which involves a further and yet elusive second state termed “m-state”. Characterizing the dynamics between these states is of paramount importance to understand the transport mechanism. The authors refolded yAAC3 from inclusion bodies in the detergent dodecylphosphocholine (DPC), and observed micro-to-millisecond (μs-ms) motion in part of yAAC3 using CPMG NMR experiments. The authors propose that this asymmetrically distributed dynamics, involving residues located in three out of six helices (α1, α2 and α6), corresponds to excursions from the c-state, which they believe is the predominant state in their sample, to an alternative state populated to 2%. They further propose that this transiently populated state might correspond to, or at least be similar to, the long-sought m-state. Support for the importance of the sparsely populated state for functional turnover comes from their finding that the dynamics is quantitatively different when the substrate (ADP) or the inhibitor (CATR) is present. As discussed below, we disagree with their interpretations. Specifically, we believe that (i) the sample was not in a functionally relevant folded state, (ii) the dynamics fits were not correctly performed and the dynamic parameters are quantitatively incorrect, and (iii) the substrate or the inhibitor have negligible effects on the dynamics in their sample, thus challenging their main conclusions.

A number of studies have demonstrated the denaturing effects of DPC on AAC and related mitochondrial carriers.^4–7^ The authors chose to study binding of CATR and ADP as criteria to assess whether their yAAC3/DPC sample was functional. They report a dissociation constant (K_d_) for CATR-binding of about 150 μM. The apparent agreement with one particular literature value^8^ would suggest that yAAC3 is functional in DPC. However, this literature value was a typographical error and has been corrected to 192 nM (instead of 192 μM)^9^, in agreement with many previously reported low-nM K_d_ values in membranes and detergents^10–13^. The CATR affinity in DPC is, therefore, ca. three orders of magnitude lower than the one for a functional protein. This finding by itself strongly suggests that the sample is in a non-native state.

Furthermore, based on NMR spectra of yAAC3 in DPC with CATR and ADP, the authors proposed that the interactions with CATR and ADP are specific. They support this claim through chemical-shift perturbation (CSP) data observed upon CATR titration for residues S31, K27, I84, R85 and T86 (reported in a very small excerpt of the NMR spectra in Figures 1e and S1 of ref. 1), and CSPs upon addition of ADP (Figure 3c of ref. 1). We have reanalyzed the titration data, using the spectra provided by the authors, and find that CSPs upon addition of CATR or ADP are spread throughout the molecule (Figure 1A). Residues with the largest ADP-induced CSPs do not point towards the central cavity, where ADP binds,^14–17^ but are on the mitochondrial-matrix side, over 20 Å away from the presumed binding site. Likewise, numerous residues experiencing CSPs with CATR lie more than 20 Å away from the CATR binding site known from crystal structures2, and are particularly located in the N-terminal loop and the matrix helices, least expected to be involved in CATR binding (Figure 1B). Possibly, unspecific electrostatic interactions in these sample conditions of low ionic strength might be an explanation for the observed effects.

**Figure 1.**
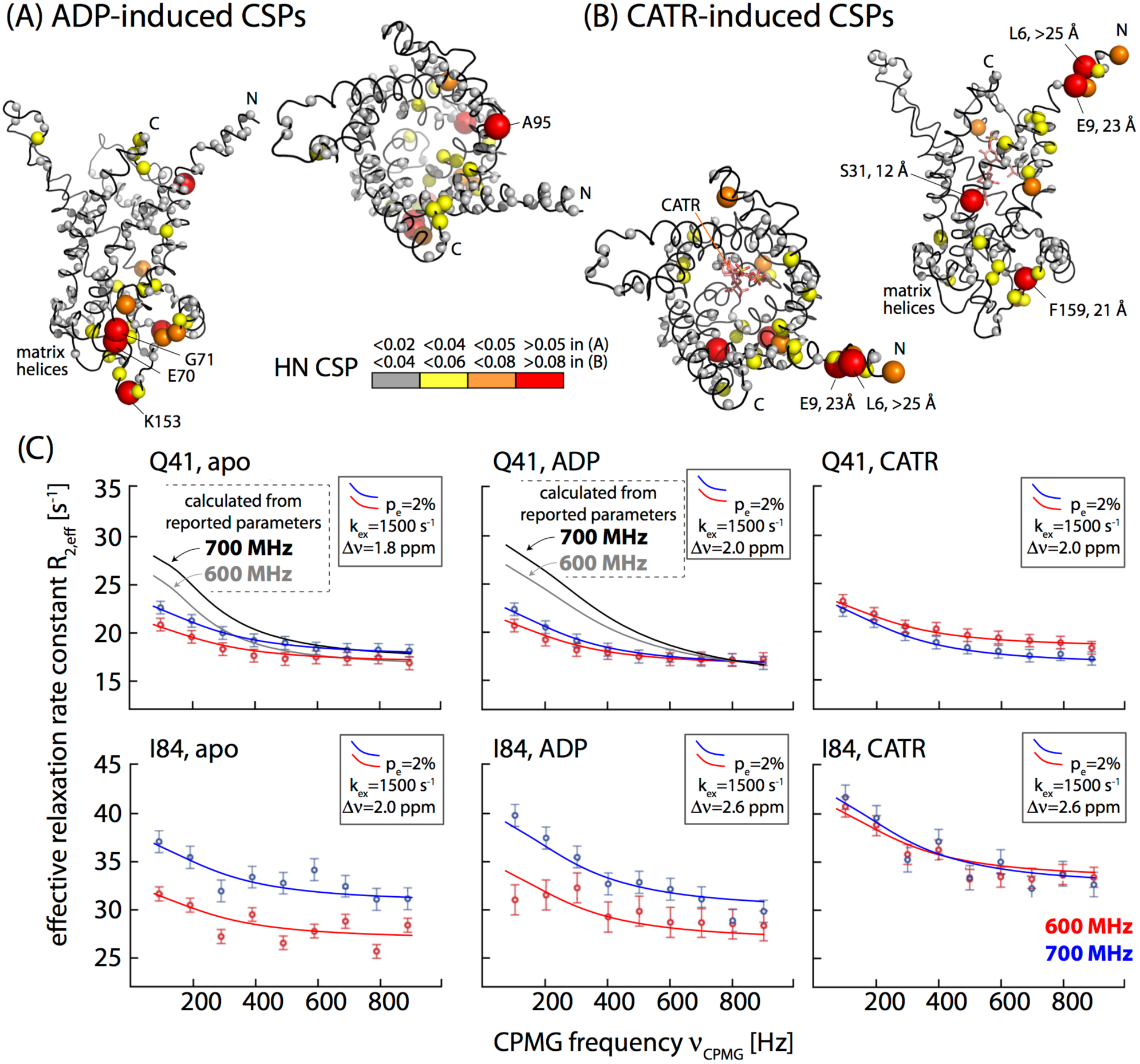
Solute-induced NMR chemical-shift perturbations interactions (A: 40 mM ATP, B: 3.5 mM CATR) and millisecond motions (C) of yAAC3 in DPC. The CSP data, plotted in (A, B) onto the structure (loops were modeled onto PDB ID 4C9J with Swiss-Model) were obtained from the original spectra of ref. 1, see Figures S2 and S3. The identity of residues with large CSPs and their distance to CATR are indicated. (C) CPMG relaxation dispersion data at two static magnetic field strengths (600, 700 MHz) for reported amide sites, Q41 and I84, in the ligand-free (left), ADP-bound (middle) and CATR-bound (right) sample conditions, as reported in Figure 2 of ref. 1 are shown as circles. Theoretical CPMG curves using the exchange parameters reported in ref. 1 are shown in grey/black in two panels (see Figure S1 for all curves). The red/blue curves were generated using a single uniform exchange rate constant k_ex_ (1500 s^-1^) and minor-state population (2%) for all residues in all three experimental conditions. Residue-wise chemical-shift differences were assumed and are reported in each panel. Note that the vertical offset of these curves is not relevant for the analysis of conformational exchange; for example, the CPMG curves shown for I84 with ADP and CATR are identical, and differ only by a constant offset. Additional data and details about the analyses of CPMG data can be found in Figure S1.

We also challenge the technical correctness and interpretation of the dynamics analyses, which form core result of the publication. We used the exchange-rate constants (k_ex_), population of “excited” state (p_e_), as well as chemical-shift differences (Δν) reported in ref. 1, to generate theoretical CPMG dispersion curves, and superimposed them onto the original CPMG data (Figure 1C, grey/black). The strong disagreement with the experimental data shows that the fit parameters by Brüschweiler *et al*. do not describe their experimental data, thus questioning an interpretation that the authors put forward: they used the large values of Δν, particularly in the proline kink regions, as an argument to propose that the excited state may resemble the m-state. Given that the actual chemical-shift changes are likely much smaller than the reported ones, this interpretation is erroneous.

We then focus on the central claim of the publication, namely that the presence of ADP or CATR significantly alters the rate at which yAAC3 exchanges with its alternate state, which goes from k_ex_= 870 ± 200 s^-1^ without substrate/inhibitor, to 1800 ± 350 s^-1^ in the presence of ADP, and 150 ± 110 s^-1^ in the presence of CATR, whereby the excited-state population (2%) remains constant. Although these absolute values are in all likelihood quantitatively wrong (see above), we investigated whether the reported CPMG data (Q41 and I84) indeed support significant differences in the dynamics upon addition of ADP or CATR. Figure 1C shows that a single set of exchange parameters (k_ex_, p_e_) for all three samples states (apo, CATR- and ADP-loaded) can describe the experimental data points similarly well as the original fit data. This analysis, although performed only with the limited data available in ref. 1, and without the 800 MHz data, which have not been made available to us, suggests that the exchange dynamics in the three different samples is likely not statistically altered by the substrate and the inhibitor. Interestingly enough, a very recent study with a non-functional mutant of yAAC3 in DPC finds essentially the same dynamics than that of WT yAAC3,^7^further supporting our view that the observed motions are not related to function.

It is unlikely that in functional AAC, CATR has only the very modest (if not negligible) effect on the dynamics, and that it leads to the modest spectral changes reported in ref. 1 (cf. Figure S3). CATR is a very strong inhibitor that locks native AAC into a single conformation. It remains bound to native yAAC3 extracted from membranes even during days of purification steps without externally added CATR.^2^ The thermal stability of AACs in mild detergent increases by more than 30°C upon binding of CATR,^5,6^ revealing a considerable change in the properties induced by inhibitor-binding, at variance with the very modest CATR-induced effects reported in ref. 1. The lack of binding specificity, low affinity and hardly detectable effects on dynamics all point to a non-functional state of yAAC3 in DPC, in line with other reports on mitochondrial carriers.^4–7^

This work was supported by the European Research Council (ERC-StG-311318).

## Supporting Information

**Figure S1 (preceding page).**
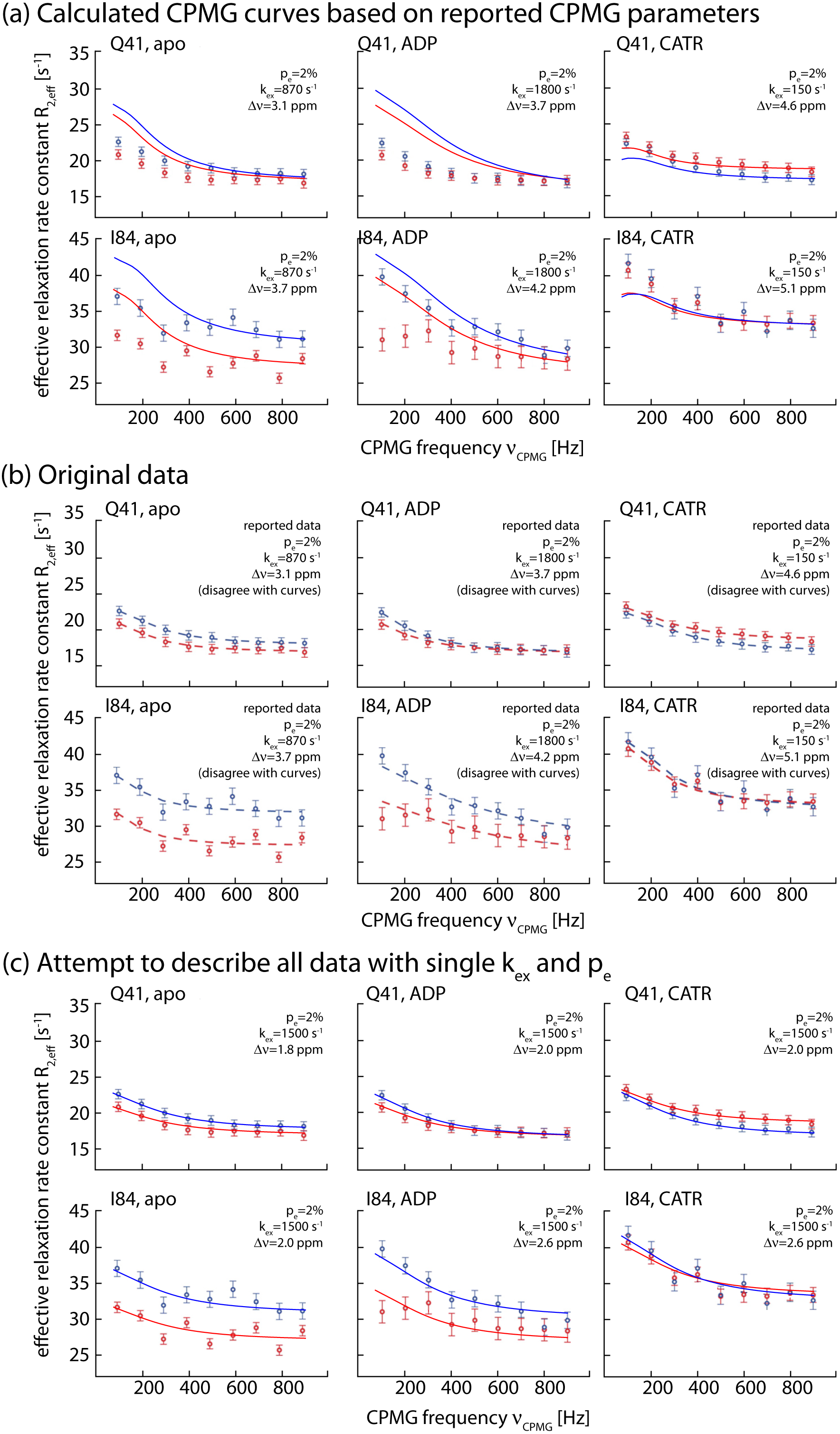
CPMG relaxation dispersion data of yAAC3 in DPC micelles, as reported by Chou and co-workers. In all cases, (a), (b) and (c), the same experimental data are shown, as provided in Figure 2 of the aforementioned publication. Herein, data shown in red and blue were obtained at ^1^H Larmor frequencies of 600 and 700 MHz, respectively. In (a) we plot theoretical CPMG curves onto these data, using the exchange parameters (minor-state population, chemical-shift difference, exchange-rate constant) reported in the publication and a numerical solution of the Bloch-McConnell equations (McConnell 1958). The exchange parameters used in these calculations are stated explicitly in each panel here. While generating these theoretical curves, we added a vertical offset 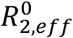, corresponding to the plateau of R_2,eff_ that is reached at high CPMG frequency. This offset, which does not bear any physical information as far as the conformational-exchange process is concerned, is an additional parameter that has to be fitted in all CPMG analyses. It was chosen here to visually match the data, and is of no direct relevance for this comparison. The comparison of the theoretical curves with the experimental data reveals that the reported parameters are in clear disagreement with the experimental data points. We have performed a similar analysis also for the 800 MHz data reported in the Supporting Information of ref. 1, and we find a similar disagreement with the theoretical curves. Panels shown in (b) are the original data of Figure 2 of the aforementioned publication, including the original relaxation dispersion curves. For completeness, we have noted the exchange parameters reported there, even though these parameters do actually not properly describe the curves plotted, as highlighted in panel (a). It is not clear at this point which exchange parameters were used to generate these curves. In panel (c) we show that the reported data do not support that the exchange dynamics is different when ADP or CATR are added. We have used in all these panels the same exchange rate constant (1500 s^-1^) and population of the minor state (2%). We have allowed two parameters to be modestly different between the three sample conditions and two different residues, namely (i) the chemical-shift difference, Δν and (ii) the vertical offset, 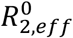 Allowing Δω to vary in the different scenarios (apo, ADP-, CATR-bound) is reasonable because the respective ground-state chemical shifts differ, and thus it is likely that the Δν may also differ. The (very modest) changes in Δω between the different samples do not imply that the dynamics differs. Likewise, adjusting the plateau level, 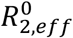, is inconsequential, because this parameter is without physical meaning in the context of exchange dynamics. The fit quality of this scenario, with substrate/inhibitor-independent dynamics is similar to the original fits, shown in panel (b). This finding suggests that the dynamics of yAAC3 in DPC micelles is not significantly altered by the presence of substrate/inhibitor, at least as far as one can judge from the data provided by Chou and co-workers.

**Figure S2.**
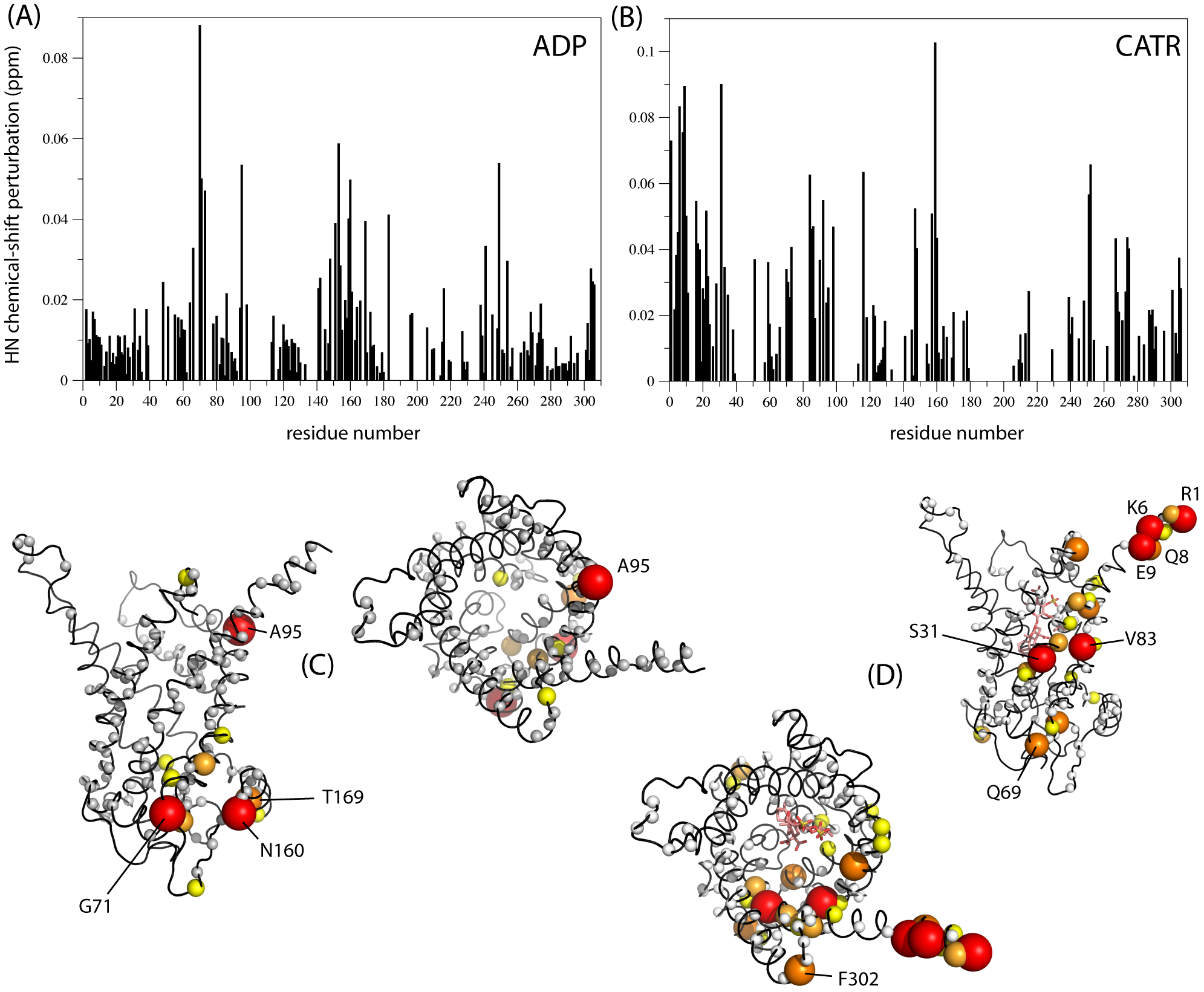
Chemical-shift perturbation upon addition of 40 mM ADP or 3.5 mM CATR to yAAC3. For generating the residue-wise plots of (A) and (B), which are the basis of Figure 1A and 1B, we have reanalyzed the original spectra from ref. 1, kindly provided by the authors. The combined ^1^H^15^N CSP was calculated as the sum of the absolute chemical-shift changes in the two dimensions, whereby the ^15^N contribution was weighted by 1/5 to account for the larger range (in ppm) of ^15^N than ^1^H. This scaling factor, often used to calculate combined CSPs (Williamson 2013), has been used also in ref. 1 for generating the data shown in C, D. For comparison, we have also plotted the CSPs reported by the authors of ref. 1 and deposited recently in the BioMagResBank (entry 27252). While there are differences in the two analyses, this data also highlights that many residues affected by the substrate and the inhibitor are located far from the binding sites, providing a qualitatively similar picture as Figure 1. Note that Arg1, one of the residues with strongest CATR-induced CSPs, is not present in the native sequence of yAAC3.

**Figure S3.**
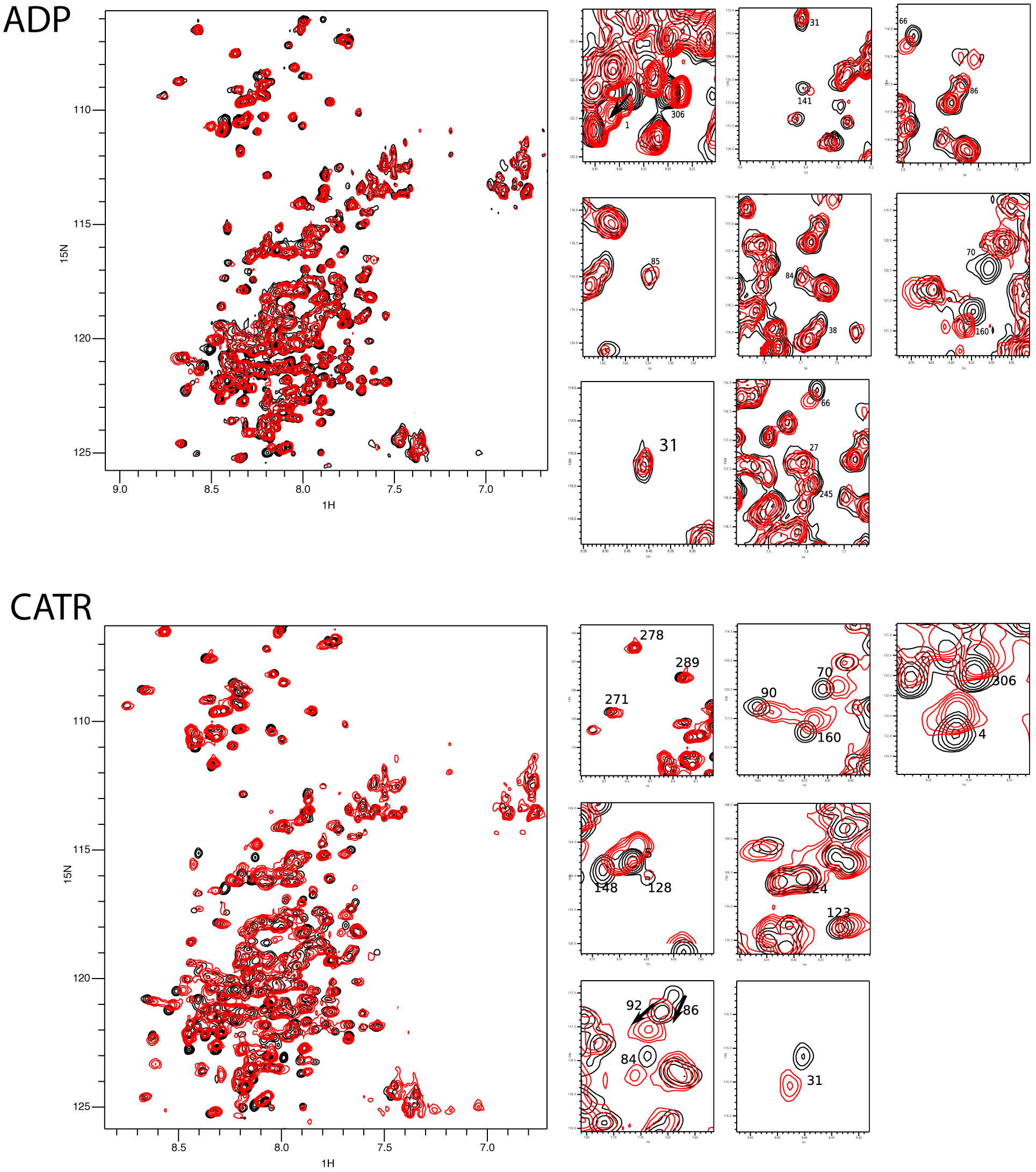
Spectra of yAAC3 in the apo state (black) and with 40 mM ADP or 3.5 mM CATR. Assignments of selected residues are shown in the zoom panels on the right. Spectra have been kindly provided by Dr. Sven Brüschweiler and Prof. J. J. Chou.

